# BLINK: Ultrafast tandem mass spectrometry cosine similarity scoring

**DOI:** 10.1101/2023.02.14.528550

**Authors:** Thomas Harwood, Daniel G.C. Treen, Mingxun Wang, Wibe de Jong, Trent R. Northen, Benjamin P. Bowen

## Abstract

**Summary:** Metabolomics has a long history of using cosine similarity to match experimental tandem mass spectra to databases for compound identification. Here we introduce the Blur-and-Link (BLINK) approach for scoring cosine similarity. BLINK calculates substantially equivalent cosine similarity scores (>99% identification agreement) over 1000 times faster than commonly used loop-based implementations by bypassing fragment alignment and simultaneously scoring all pairs of spectra using sparse matrix operations. This performance improvement can enable calculations to be performed that would typically be limited by time and available computational resources.

**Availability and Implementation:** BLINK is implemented in Python3 and is published under a modified open source license. Code and license are available on Github: https://github.com/biorack/blink

## 1 Introduction

Tandem mass spectrometry has become a critical component of both targeted and untargeted metabolomics experiments. Comparing fragmentation spectra between authentic standards and experimental data is central to making high confidence assignments in targeted metabolomics experiments. Untargeted experiments depend on matching experimental MS2 spectra against databases such as GNPS[1], Metlin[2], and MassBank[3]. Although different approaches have been developed[4,5], this is still conventionally done by aligning fragment ions that share the same mass-to-charge ratio (*m/z*) and calculating the cosine similarity of their intensities. As a consequence of early mass spectrometers having low resolution, integer binning was a sufficient alignment step for comparing fragmentation spectra[6]. But as the resolution of mass spectrometers grew, so too did the sophistication of alignment algorithms necessary for high confidence identification and similarity scoring. Many of these approaches use iterative operations to compare individual fragment ions between reference and experimental spectra, which is inherently slower than vectorized operations.

To this end we introduce BLINK, an approach that enables scoring of fragmentation spectra using sparse matrix operations without the drawbacks of traditional *m/z* binning based approaches. Rather than multiplying two matrices of binned fragmentation spectra directly, we use a uniform kernel to link together bins that are within machine noise tolerance, thereby vectorizing the previously costly alignment step. Matching ion counts can be approximated via the same methodology. Here we describe BLINK’s implementation and present a comparison between BLINK and the CosineGreedy scoring algorithm used in MatchMS [7].

## 2 Results and Discussion

For this study, we focused our comparison on MatchMS because it is a high-performance, widely used, and well-supported Python package with overlapping use cases with BLINK. To assess BLINK’s agreement with canonical cosine-based scoring approaches, we used tandem mass spectra sampled from an aggregate of all publicly available GNPS libraries. Using both MatchMS and BLINK, two randomly selected sets of MS/MS spectra were scored against each other. Spectra were classified as “similar” if their cosine score was ≥ 0.7 and matching ions were ≥ 6. All 1829 spectra classified as similar by MatchMS were also similar using BLINK, however, 13 spectra were only classified as similar by BLINK (Fig. 1a). The spectra classified as similar by both BLINK and MatchMS had raw scores that varied by a mean value of 0.0004 and matching ion counts that varied by a mean value of 0.06. The deviation in scores and counts is due to BLINK’s alignment-free approach, which factors all ions within tolerance into the scoring and counting, rather than selecting only one.

**Figure 1:**
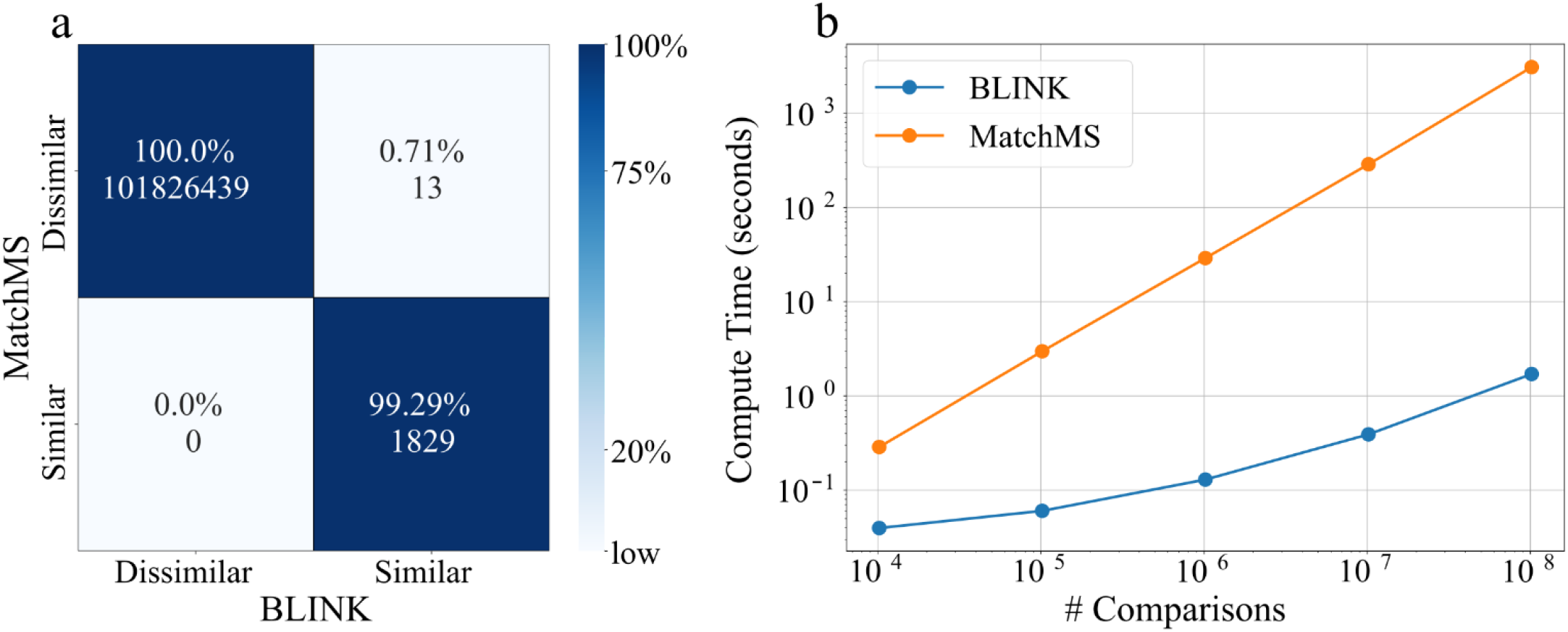
Evaluation of timing and agreement comparing BLINK cosine scoring to MatchMS. **(a)** Modified confusion-matrix of spectral similarity using BLINK and MatchMS on over 100 million comparisons where “similar” results had high scores and high matching ions. In addition to raw counts, the percentages of counts normalized by MatchMS values are shown. **(b)** Median cosine scoring runtimes for BLINK (blue) and MatchMS (orange) across 3 replicates.

While the results were similar, BLINK was much faster. Importantly, BLINK scales more favorably with high numbers of comparisons. BLINK can perform 1e8 comparisons 1806 times faster than MatchMS (1.7 seconds for BLINK vs 51 minutes for MatchMS) (Fig. 1b). This difference is because standard tools such as MatchMS rely on loop-based, pairwise alignment procedures for matching ions within machine noise tolerance during scoring. BLINK circumvents this computationally expensive fragment alignment step. In addition, rather than iteratively scoring pairs of spectra, BLINK simultaneously scores all pairs of spectra via sparse matrix operations. This speedup can enable investigators to perform high-throughput calculations that would otherwise be prohibitively time intensive. For example, NIST20[8] currently has over 1.3 million spectra and a typical metabolomics data set can have 10,000 features. Extrapolating from the speed tests in Fig 1b, performing a complete database search would take 4.5 days with MatchMS, and this is reduced to 3.3 minutes using BLINK.

There are some limitations to BLINK’s alignment-free approach. Because all ions within tolerance are factored into the cosine score, scores calculated using BLINK diverge from precise, loop-based implementations as tolerance increases. Therefore, traditional tools may be a more appropriate choice for data generated by low resolution mass spectrometers that require a wide tolerance window. However, given trends in mass spectrometry hardware development, this limitation will likely become less relevant over time as higher resolution machines are continually developed[9]. We envision that the speed-up using BLINK will enable scientists to efficiently make chemical assignments as the size of MS2 databases continues to grow, particularly *in-silico* databases uncoupled from the availability of new authentic standards and instrument throughput [10,11].

## 3 Implementation

Typically two fragmentation data files, a query and a reference, are used as input. The files are parsed and the fragmentation spectra are represented as lists of *m/z* and intensity arrays with associated metadata. To discretize the spectra, *m/z* values are first converted to integer-bins based on the user defined bin width (default is 0.001 Da). Intensity arrays in each spectrum are then unit-vector normalized. Each set of processed spectra is converted into two sparse matrices. The first contains fragment intensities, and the other fragment counts. Each sparse matrix is constructed such that rows are *m/z* bins and columns are the spectrum index.

Prior to scoring, the *m/z* bins are expanded (i.e. blurred) by distributing intensities and summing fragment *m/z* bins with a tolerance-window wide array, referred to here as BLINK’s kernel. This allows for *m/z* bins to be linked between spectra within a tolerance parameter (default is 0.01 Da). It is only necessary to perform this computation on one set of spectra, so the kernel is used to transform the smaller of the two (typically the query).

Each pair of sparse matrices are resized such that they contain an equal number of rows, and one of the matrices is transposed. The two matrices are multiplied, generating the score matrix. When performed on the matrices containing intensity data, this step simultaneously computes the cosine scores for each combination of spectra. Fragment counting is based on the same algorithm, but uses the fragment count matrices as input.

## 4 Methods

### 4.1 Speed Benchmarking & Similarity Agreement

Calculations were done using an exclusive CPU node on the Perlmutter supercomputer at NERSC. Each CPU node is equipped with 512 Gb of DDR4 RAM and dual AMD EPYC 7763 64x core processors. Two sets of spectra were randomly sampled from an aggregate of all GNPS library spectra (All-GNPS) as of January 17th 2022 ranging in size from 1e2 to 1e4. Each subset of spectra was sampled independently without replacement. Scoring was performed using tolerance values of 0.01 Da and 0.099 Da for BLINK and MatchMS respectively. The BLINK bin width parameter was set to 0.001 Da. BLINK’s true tolerance is equivalent to the tolerance parameter subtracted by the bin width, hence the difference in tolerance values used for the two algorithms. To improve scoring behavior of both MatchMS and BLINK, the spectra were filtered to remove noise ions and the intensity values were scaled by their square-root. Fragment ion noise filtering was accomplished by removing ions that were <1% of base peak intensity and ions with *m/z* values greater than the precursor *m/z*. Additionally, all ions with intensity values of 0 were removed from the spectra.

Progressively larger sets of spectra were sampled and scored with 3 replicates and their median calculation timings with MatchMS and BLINK were reported in Figure 1b. The modified confusion-matrix reported in Figure 1a was generated using 1e8 scores from the first replicate of the speed benchmark. Spectra were classified as similar if their score was ≥ 0.7 and matching ions were ≥ 6. These values were chosen because they are the GNPS default filters.

### 4.2 Mean Score & Matching Ion Count Differences

The same set of scores used to construct the confusion-matrix were used to calculate the mean score and matching ion count differences. Differences of the true positive scores (classified as similar by both MatchMS and BLINK) were calculated by subtracting the BLINK scores and counts by their MatchMS counterpart.

## 5 Conclusion

Comparison of fragmentation spectra has become a primary step in both targeted and untargeted metabolomics workflows. Using an alignment-free and vectorized approach, BLINK is able to compute cosine similarity scores of tandem mass spectrometry data faster than previously possible.

## 6 Acknowledgements

We thank Yu-Hang Tang for sharing their expertise in machine learning and linear algebra. This work was supported by the Machine Learning for Chemistry Laboratory Directed Research and Development (LDRD) Program of Lawrence Berkeley National Laboratory and the Office of Science of the U.S. Department of Energy by ENIGMA-Ecosystems and Networks Integrated with Genes and Molecular Assemblies (http://enigma.lbl.gov), a Science Focus Area Program at Lawrence Berkeley National Laboratory; the U.S. Department of Energy Joint Genome Institute (https://ror.org/04xm1d337), a DOE Office of Science User Facility; and the National Energy Research Scientific Computing Center (NERSC) a DOE Office of Science User Facility all operated under Contract No. DE-AC02-05CH11231 with the U.S. Department of Energy. The U.S. Government retains, and the publisher, by accepting the article for publication, acknowledges, that the U.S. Government retains a non-exclusive, paid-up, irrevocable, world-wide license to publish or reproduce the published form of this manuscript, or allow others to do so, for U.S. Government purposes.

